# Understanding the Athena SWAN award scheme for gender equality as a complex social intervention in a complex system: analysis of Silver award action plans in a comparative European perspective

**DOI:** 10.1101/555482

**Authors:** Evanthia Kalpazidou Schmidt, Pavel V. Ovseiko, Lorna R Henderson, Vasiliki Kiparoglou

**Affiliations:** Department of Political Science, Aarhus University, Denmark; Radcliffe Department of Medicine, Medical Science Division, University of Oxford, Oxford, United Kingdom; NIHR Oxford Biomedical Research Centre, Oxford, United Kingdom; Nuffield Department of Primary Care Health Sciences, Oxford, United Kingdom

**Keywords:** Athena SWAN, gender equality award scheme, Medical Sciences, complexity approach, typology of interventions, European comparative, responsible research and innovation

## Abstract

**Background:** Given that the complex mix of structural, cultural, and institutional factors has produced barriers for women in science, an equally complex intervention is required to understand and address them. The Athena SWAN award scheme for gender equality has become a widespread means to address barriers for women’s advancement and leadership in the United Kingdom, Ireland, Australia, the United States of America, and Canada, while he European Commission is exploring the introduction of a similar award scheme across Europe.

**Methods:** This study analyses the design and implementation of 16 departmental Athena SWAN Silver action plans in Medical Sciences at one of the world’s leading universities in Oxford, United Kingdom. Data pertaining to the design and implementation of gender equality interventions were extracted from the action plans, analysed thematically, coded using categories from the 2015 Athena SWAN Charter Awards Handbook, and synthesised against a typology of gender equality interventions in the European Research Area. The results were further analysed against the complexity research literature framework, where research organisations are perceived as dynamic systems that adapt, interact and co-evolve with other systems.

**Results:** Athena SWAN is a complex contextually-embedded system of action planning within the context of universities. It depends on a multitude of contextual variables that relate in complex, non-linear ways, and dynamically adapt to constantly moving targets and new emergent conditions. Athena SWAN Silver action plans conform to the key considerations of complexity: 1) multiple actions and areas of intervention with a focus on the complex system embedded in and the local dynamics, 2) the non-linearity of interventions and the constantly emerging conditions, 3) impact in terms of contribution to change, improved conditions to foster change, and the increased probability that change can occur.

**Conclusions:** To enact effective sustainable gender equality structural and cultural change, it is necessary to acknowledge and operationalize complexity as a frame of reference. Athena SWAN is the single most comprehensive and systemic gender equality scheme in Europe and can be strengthened further by promoting the integration of sex and gender analysis in research and education. Gender equality policies in the wider European Research Area can benefit from exploring Athena SWAN’s contextually-embedded systemic approach to dynamic action planning and inclusive focus on all genders and categories of staff and students.

## Background

Given that the complex mix of structural, cultural, and institutional factors produces barriers for women in science, an equally complex intervention is required to address them [1]. It has been argued that gender inequalities persist due to “culture, processes and practices that constitute the structural systems of contemporary organizations and therefore are taken for granted and mostly left unchallenged” [2–4]. Several studies also reveal how deep-rooted assumptions about academia being a meritocracy reproduce inequalities, and call for scrutinizing the existing practices, procedures and structures [5–8]. While the structure of academic organizations reproduces gender stereotypes, such as power relations and distribution of women and men at top level positions, career prospects, etc. [9–11], policy makers and academic leaders have for a long time failed to recognize and address institutionalized structural and cultural barriers that hinder women’s advancement and leadership in academic and research organizations [5]. In the recent decade, the focus on structural and cultural barriers has gained prominence and been successively “built into the funding schemes of different agencies supporting interventions to reduce gender inequality in science” [12]. Most notably, the Athena SWAN Charter globally and gender equality policies in the European Research Area provide support and incentives for academic and research organizations to address structural and cultural barriers to women’s advancement and leadership through action planning.

The Athena SWAN Charter was established in the UK in 2005 to encourage and recognize the commitment of higher education and research institutions to advancing careers of women in science, and in 2015, it was expanded to arts, humanities, social sciences, business, and law [13]. When institutions join the Athena SWAN Charter, they commit to systematically assessing and advancing gender equality through action planning and apply for awards recognizing their success. Applications are peer reviewed by academics, subject experts, human resources and equality and diversity practitioners from other member institutions who then make recommendations on the level of awards [13]:

- A Bronze award requires an assessment of gender equality and the related challenges as well as a 4-year action plan to address these challenges;
- A Silver award recognizes the successful implementation of the proposed action plan and its measurable impact;
- A Gold award recognizes beacons of achievement in gender equality and champions in promoting good practice in wider community.

Athena SWAN has become a common means to address barriers for women’s advancement and leadership in the UK, Ireland, and Australia; the United States of America (USA) and Canada use modified approaches, and discussions are underway in India and Japan. In the UK, the National Institute for Health Research (NIHR) requires the Athena SWAN Silver award as a prerequisite for applying for competitive biomedical research funding [14]. In Ireland, by 2023, higher education and research institutions will be required to hold the Athena SWAN Silver award to be eligible for competitive government research funding [15]. In Australia, the Australian Academy of Science and the Australian Academy of Technology and Engineering runs the Athena SWAN gender equality award scheme as part of the Science in Australia Gender Equity (SAGE) program [16]. In the USA, the American Association for the Advancement of Science (AAAS) uses a modified Athena SWAN self-assessment and improvement framework as part of its STEM Equity Achievement (SEA) Change programme [17]. In Canada, the government has committed to implementing a “made-in-Canada” Athena SWAN initiative [18]. India and Japan have also expressed interest in trialling the Athena SWAN framework [19].

In the wider European Research Area (ERA), three strategic objectives for fostering gender equality in research and innovation are promoted: (i) more women in research, (ii) more women in leadership positions, and (iii) the integration of the gender dimension in research and curricula [20]. The development and implementation of gender equality action plans is a key instrument for promoting structural and cultural change in the European Research Area. Although the European Commission (EC) adopted the structural change approach as late as in 2011 [21], efforts have significantly intensified in recent years, particularly, following the introduction of the Responsible Research and Innovation approach [22].^1^ A number of multi-national European projects focus on the implementation of gender equality action plans tailored to research institutions in a number of European countries producing a substantial knowledge base and establishing a set of best practices. Yet, in some countries the development of GE action plans still remains in an embryonic stage, and in some other countries where implementation is underway, institutional change needs to be further promoted and evaluated [23, 24].^2^ To further activate cultural and structural change in research organisations and universities wide across Europe, the European Commission explores scenarios for the introduction of a gender equity award scheme similar to Athena SWAN [25].

In this paper, based on the most recent research [12, 26] and rich empirical data, we argue that to activate effective gender equality structural and cultural change, it is necessary that interventions acknowledge and operationalize the notion of complexity as their frame of reference. The paper is organized as follows. We first present our methods. Secondly, we focus on the design and implementation of 16 Athena SWAN Silver action plans in Medical Sciences at one of Europe’s leading universities, Oxford University, analyse gender equality interventions thematically, and present the results in a comparative European perspective using the complexity approach. Finally, we discuss Athena SWAN as a complex social intervention, formulate practical implications for impact assessment of Athena SWAN and other complex gender equality interventions, and outline strategic opportunities to strengthen gender equality policies in the European Research Area.

## Methods

### Study aim and objectives

The aim of this study is to analyse Athena SWAN Silver action plans in Medical Sciences at Oxford University, using a complexity approach and a comparative European perspective. The objectives of the study are as follows:

- To explore what types of interventions are associated with Athena SWAN Silver action plans.
- To compare these types of interventions with the types of interventions used in the wider European Research Area.
- To discuss how a complexity approach can provide insights for policy and practice.

### Study setting and context

The study is set within the context of Medical Sciences at the University of Oxford. The university tops the Times Higher World University Rankings [27], outperforms other universities in EU research funding competitions [28], and files more international patent applications to the World Intellectual Property Organization than any other university in Europe [29]. To ensure its competitive advantage, the university is actively seeking new ways to attract the best students, recruit and retain the most talented staff, and increase research funding. Thus, advancing gender equality in education, research, and innovation is one of the key strategic priorities for the university [14, 30–32]. It has been a member of the Athena SWAN Charter since its establishment in 2005 and, as of October 2018, holds 19 Silver and 15 Bronze departmental level awards in Medical Sciences, Mathematics, Physics, Life Sciences, and Social Sciences. The overwhelming majority (16/19) of Silver awards are in Medical Sciences, reflecting the linkage of Athena SWAN Silver awards to NIHR competitive biomedical research funding.

### Data collection and analysis

The study is based on the document analysis of all 16 departmental Athena SWAN Silver action plans in Medical Sciences at the University of Oxford. We collected the required action plans from a dedicated university webpage, all successful award applications and action plans are publicly available [33]. We extracted data pertinent to the design and implementation of gender equity interventions, classified the target population of each action by gender and staff category, analysed these data thematically [34], and coded the emerging themes in Microsoft® Office using categories from the 2015 Athena SWAN Charter Awards Handbook [35]. In doing so, we were sensitised by the concept of complexity and used the process of constant comparison to synthesise emerging themes in a comparative European perspective, as described below. We reached consensus on themes and comparison by agreement. We also reflected on our own prior views and experiences, which may have influenced our analysis and interpretation of data.

### Complexity approach

Under the complexity approach [36, 37], interventions are considered as part of the complex system within which they take place. A complex system is characterised by a multitude of components, which through continuous interaction create a system-level organisation with the whole greater than the sum of its components and by positive feedback processes, in which outcomes of the process are necessary for the process itself [38]. Typically, a complex system operates in a multi-layered context and dynamically adapt in response to changes in the environment. In complex systems there is a multitude of contextual variables that interact in complex, nonlinear ways [36]. “Complex systems are adaptive—they respond to changes. This central feature of complex systems is what makes them distinct from systems that are merely complicated” [36].

Complexity in research organisations is a dynamic complex framework that adapts, interacts and co-evolves with other systems, rather than being a stable arrangement of different contextual features [26, 39, 40]. What embodies complexity in complex interventions is disproportionate interactions (e.g. a proportionally small intervention can make a vast difference in some point in time), recursive causality with strengthening loops, and emergent outcomes that need to be addressed instantaneously [37]. Greenhalgh and Papoutsi [40] conclude that instead of a linear, cause-and-effect causality, the complexity paradigm is characterized by “emergent causality: multiple interacting influences account for a particular outcome but none can be said to have a fixed ‘effect size’”, and a pragmatic adaptation to changing contexts and emerging circumstances. Another aspect of complexity that helps us understand and manage variations across local contexts is self-organisation [41]. Manifestation of self-organisation takes place through specific action patterns that depend on locally available resources and local contextual conditions that can promote or impede successful implementations [42]. Operationalizing the notion of complexity as a frame of reference implies thus multiple areas of intervention with a focus on the local dynamics [26].

### Comparative European perspective

We analyse Athena SWAN interventions in a comparative European perspective based on a typology of gender equality interventions in research and innovation developed within the EU Horizon 2020 project EFFORTI (Evaluation Framework for Promoting Gender Equality in Research and Innovation) [43] Initially proposed by Kalpazidou Schmidt and Cacace [26] based on an empirically study of 109 GE interventions worldwide and a literature review [43], the EFFORTI typology of gender equality interventions in research and innovation was subsequently adapted and expanded to synthesise knowledge on gender equality interventions from major European projects, such as GEAR (Gender Equality in Academia and Research) [24], GEDII (Gender Diversity Impact – Improving research and innovation through gender diversity) [44], GENERA (Gender Equality Network in the European Research Area) [45], Gender-NET (Promoting Gender Equality in Research Institutions and Integration of the Gender Dimension in Research Content) [46], PRAGES (Practicing Gender Equality in Science) [47], and STAGES (Structural Transformation to Achieve Gender Equality in Science)[48]^3^. An overview of the EFFORTI typology of gender equality interventions in research and innovation is presented in Additional file 1.

## Results

In total, 16 Athena SWAN Silver action plans contain 547 actions pertaining to gender equality interventions. On average, there are 34 actions per action plan. The target population of the Athena SWAN Silver actions analysed by gender are all genders indiscriminately (88%), women (11%), and men (1%). The target population of the Athena SWAN Silver actions analysed by student and staff category are academic and research staff (52%), all staff (32%), students (1%), students and staff (4%), and professional and support staff (1%). Thematic analysis of actions against the relevant sections and sub-sections of the 2015 Athena SWAN Charter Awards Handbook [35] has resulted in five themes and 22 sub-themes. According to frequency analysis of actions by theme, “Organisation and culture” (28%) and “Career development” (28%) are the most frequent themes. They are followed by “Self-assessment and monitoring” (17%), “Key career transition points” (15%), and “Flexible working and career breaks” (13%). Themes and sub-themes are summarised and visually represented in Figure 1 using the sunburst chart technique whereby colour-coded concentric circles display a hierarchical relationship between major themes and sub-themes in proportion to the frequency of actions pertaining to each theme and sub-theme.

**Fig. 1.**
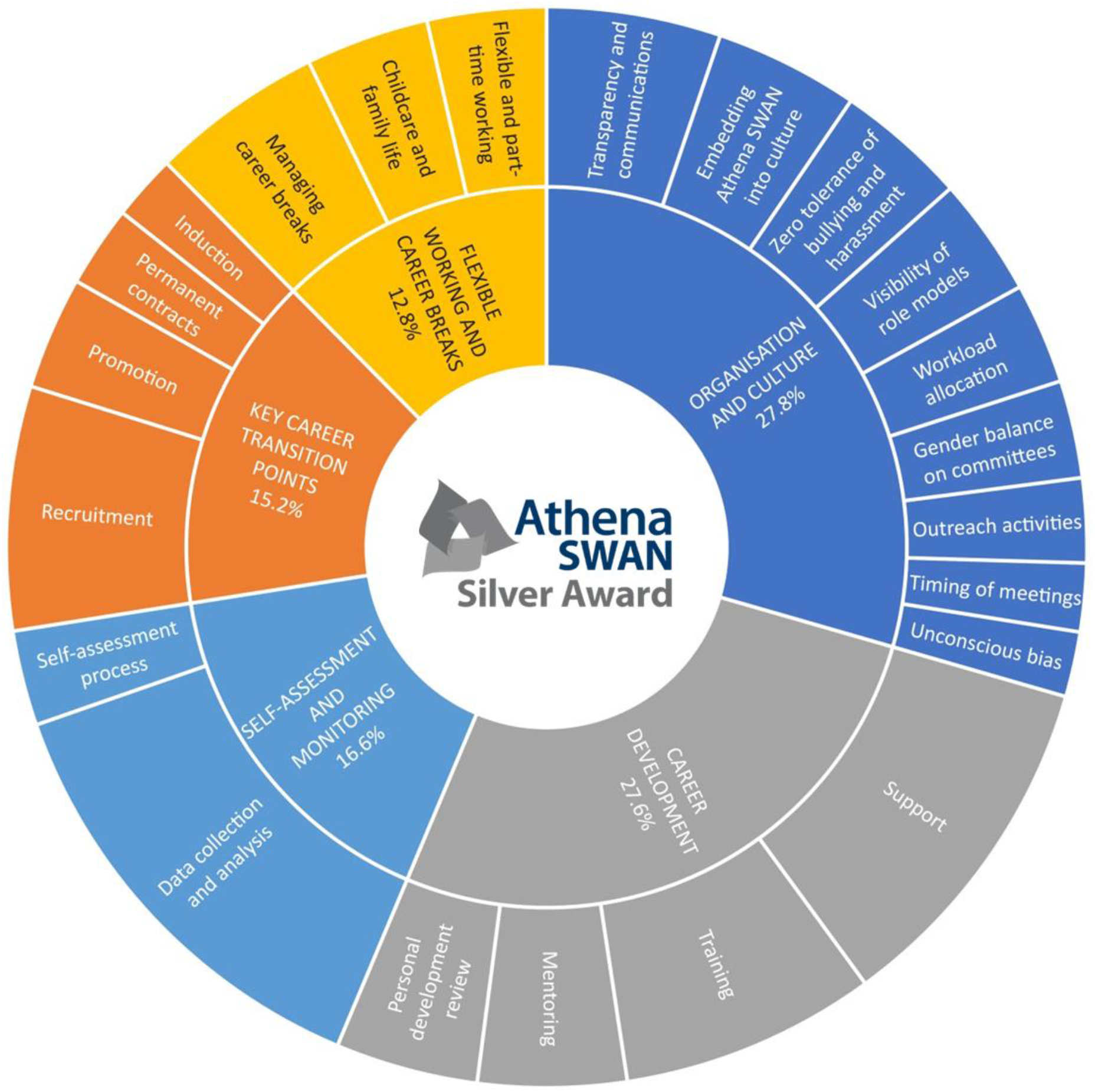
Athena SWAN Silver award interventions by theme, sub-theme, and frequency of actions in 16 departmental action plans in Medical Sciences at the University of Oxford, 2014-2017.

A comparison of the SWAN Silver action plans with the EFFORTI typology of gender equality interventions in research and innovation reveals that they represent 78% (31/40) of gender equality interventions types used in the wider European Research Area and a further 8 distinctive intervention types (Table 1). Gender equality interventions in the wider European Research Area tend to target primarily women in academic and research roles to address their underrepresentation in European research organisations through a range of interventions, including among others affirmative action such as quotas, funding and positions reserved to women. Athena SWAN has a somewhat broader focus as regards the target population, including also professional and support staff and students as well as considering the intersectionality of gender and other aspects of identity, such as sexuality, race, disability, age, and religion. Yet, Athena SWAN lacks the intervention types based on the wider ERA objective of integrating the gender dimension in research and education (see Table 1, gender equality interventions distinctive to EFFORTI points 5-9).

**Table 1.** Comparison of gender equality interventions distinctive to Athena SWAN and EFFORTI.

| Gender equality interventions distinctive to Athena SWAN | Gender equality interventions distinctive to EFFORTI |
| --- | --- |
| 1. Institutionalised self-assessment teams<br>2. Intersectional approaches to data collection and analysis<br>3. Addressing a full continuum of key career transition points<br>4. Career development interventions targeting professional and support staff<br>5. Career development interventions targeting students<br>6. Mandatory training on unconscious bias and bullying and harassment<br>7. Revising timing of meetings and events<br>8. Workload allocation model | 1. Institution of quotas<br>2. Introduction of chairs and positions reserved to women<br>3. Special funding for women researchers<br>4. Support to mobility, including spouse relocation schemes<br>5. Integration of the gender dimension and impact in research<br>6. Integrating the gender dimension in tertiary education<br>7. Revision of teaching curricula and texts<br>8. Introduction of single-sex degree and specialisation courses<br>9. Provision of gender and women's studies or modules |

In what follows, themes and sub-themes of the analysed actions are presented in the order of apperance in the Athena SWAN Silver award application, together with 93 illustrative examples of actions demonstrating the range of actions in each sub-theme. Illustrative actions are presented together with the code corresponding to the name of the department and the numbering of actions in the relevant action plan. The names of departments and their codes are presented in Additional file 2.

### Self-assessment and monitoring

#### Self-assessment process

To participate in the Athena SWAN Charter, every department has established a self-assessment team (SAT) with broad representation of people and skills across the department and the remit to reflect on quantitative and qualitative data, evaluate relevant policies and practices, establish priority areas and targets, develop an evidence-based action plan, and evaluate its effectiveness against the agreed objectives [35]. Specific actions regarding self-assessment focus on institutionalising the SAT and its working groups within the departmental structure and securing necessary leadership and administrative support:

- “The SAT will develop and publish terms of reference including guidelines on purpose, recruitment to the committee, roles, length of service and will embrace a vision of committee aims.” (D1, 2)
- “Embed work of the Athena SWAN Career Development Working Group in permanent training and development infrastructure of the [department].” (D16, 6.3)
- “SAT will meet termly to discuss the implementation and progress of the Silver action plan” (D15, 1)
- “Explore appointment of a new post, an Athena SWAN lead, to the administrative team.” (D8, 1.1)

#### Data collection and analysis

Departmental SATs use a variety of sources, including surveys, focus groups, interviews, and databases to collect and analyse data on gender equality among staff and students at all levels. Increasingly, SATs employ intersectional approaches to better understand the issues at the intersection of gender and other aspects of identity, such as sexuality, race, disability, age, and religion [50]. Actions to improve data collection and analysis range from making it more regular, reaching out to broader student and staff populations, and refining existing questions to trialling new methods, carrying out new types of analysis, and investigating new questions:

- “The SAT will run regular staff/student surveys and convene focus groups to assess the impact of our action plan” (D4, S1.2)
- “The question to determine gender in the survey should be posed as: ‘Female, male, self-defined, or prefer not to say’ to capture the full spectrum of gender identities.” (D5, 1.4)
- “Trial exit interviews to determine if they are an efficient way of capturing necessary information (balance of staff time vs. quality of information gathered).” (D5, 2.4)
- “Carry out a pay audit of all staff by grade scale point and gender.” (D13, 6.9)
- “Investigate the barriers for appointment to senior clinical posts overseas.” (D10, 4.4)

### Key career transition points

#### Recruitment

Departments strive to improve their recruitment practices by developing workforce intelligence and planning, improving the attractiveness of job opportunities to female applicants, using targeted recruitment, as well as by minimising selection bias through mandatory training, gender balance on selection panels, and more inclusive decision-making:

- “Investigate recruitment and subsequent working experience of members of minority groups working in the [department].” (D16, 2.2)
- “Identify skills shortages and underrepresentation in [research] groups; establish future staffing requirements and succession plans; present to [the Equality Committee] as a ‘Workforce Plan’.” (D12, 1.1)
- “Encourage more female applicants by highlighting that we will provide assistance when applying for nursery/childcare/school places.” (D4, S3.4)
- “Implement the new Electoral Board process for appointment of Statutory Professors and introduce equivalent [departmental] process for all other senior posts, and pilot the use of head-hunters.” (D13, 1.2)
- “Ensure that all people involved in recruiting (not just panel chairs) have completed recruitment and selection training.” (D5, 3.3)

#### Induction

Departments work to enhance the quality of their induction programmes, dedicated webpages, factsheets, and other materials for new recruits by tailoring them to different career stages, sites, and research groups, introducing networking and peer-support schemes, as well as monitor their effectiveness and check the awareness of key policies, resources, and career development opportunities:

- “Develop a tailored induction programme for senior researchers, group leaders and line managers with emphasis on line management responsibilities and the department’s family-friendly culture.” (D13, 2.1)
- “Develop with the newly formed Postdoctoral Society a ‘Buddying’ support system for new postdocs joining the department” (D4, S4.3)
- “HR to hold an induction/probationary meeting 3 months after the start date to ensure that the new starter is feeling settled and to check they are aware of policies, postdoc events, etc.” (D5, 3.7)
- “Ensure initial career development discussions are held during probation.” (D10, 3.1)
- “Continue to monitor the effectiveness of the induction process, and identify areas for improvement. Develop a mechanism to monitor the effectiveness of site-specific inductions.” (D15, 10).

#### Promotion

Departmental action plans are in place to accelerate career advancement of all eligible staff through the existing regrading, recognition of distinction, and award schemes. A range of actions includes raising awareness and transparency of promotion opportunities, conducting gender-sensitive review of promotion criteria and salaries, identifying and encouraging all eligible candidates and women specifically to apply for promotion:

- “For non-clinical academics, transparency of pay rises, promotions process and equivalency of tenure tracks need to be continuously developed.” (D9, S8).
- “Annual review of salaries to ensure parity and gender balance.” (D8, 3.4)
- “Continue to promote and develop criteria for prizes to ensure they are achievable for both men and women; actively show how women have met the criteria and provide case studies.” (D10, 2.5)
- “An annual audit will be conducted via [a university publications management system], of peer reviewed publications first authored by [early and mid-career researchers], taking account of part time work and family leave.” (D11, 2.3)
- “Continue to identify women and provide administrative support for promotion applications through [Recognition of Distinction] award for Professorships, Associate Professorships and University Research Lecturer scheme.” (D12 1.5)

#### Permanent and long-term contracts

Athena SWAN has spurred departments in Medical Sciences to provide more job security for academic and research staff, the overwhelming majority of whom compete for research funding in tough market and remain on short fixed-term contracts. Departments take action to transfer eligible staff on to open-ended contracts with support for at least as long as external research funding is available and set targets to increase the number of staff especially women on and permanent contracts:

- “Implement a transparent Department wide policy to review all staff on fixed-term contracts on a regular basis. Move staff from fixed-term to open contracts, where possible.” (D8, 3.2)
- “We will introduce a clear and transparent system to allow the transfer of senior research fellows on to permanent contracts.” (D4, S5.1)
- “Increase the proportion of Associate Professor and Full Professorial posts that are held by women from the current 38.7% (12/31) to 50% by 2018.” (D2, 5.3)
- “Investigate mechanisms underlying high attrition rate of female academic clinicians.” (D14, 3.5)

### Career development

#### Training

Departments seek to promote career development of not only academic and research staff, but also professional and support staff and students through training, including courses specifically designed for women. Actions are in place to better identify training needs of particular groups of staff and provide more in-house and external training with regards to management, leadership and negotiation skills, career planning, and grant writing:

- “Encourage management training (appraisals, project management, coaching, time management, and workload planning) for Principal Investigators, supervisors and line managers.” (D10, 3.2)
- “Organise targeted ‘How to’ workshops designed to help staff at the key career transition points (e.g. writing a grant application).” (D13, 1.9)
- “Identify senior staff, and those approaching senior grades, who are seeking training in ‘leadership’ from the [personal development review] discussion and provide 5 [departmental] funded places (up to £5,000 per place) on a leadership course for senior women.” (D8, 4.4)
- “Organise a ‘Manage Your Supervisor’ training session during May for first year DPhil students.” (D13, 5.1)

#### Personal development review

Athena SWAN has been instrumental to the institutionalisation of annual personal development review (PDR), which enables staff to have open conversations with their reviewer about their role, career aspirations, and development opportunities. Departments work to increase the awareness and uptake of PDR, provide training and guidance for reviewers and reviewees, and improve its effectiveness, especially, for researchers on a succession of short-term contracts, postdocs approaching independence, and other staff for whom PDR is particularly beneficial.

- “From 2017 we will move to undertaking PDR annually in April/May for all staff. We will provide flexibility for clinicians who would prefer a different time of year to enable their University PDR to inform their [National Health Service] appraisal.” (D11, 9.1)
- “Continue to reinforce to staff the necessity to follow-up on the PDR discussions periodically during the year in order to ensure progress towards the training and personal development goals.” (D12, 2.7)
- “Annual PDR workshops for staff, which covers the purpose of PDR, guidance on the conduct of PDR and the rationale for why the Department aims to increase uptake.” (D8, 4.2)
- “Add a checklist to the PDR form to encourage discussion of scientific engagement, internal and external mentoring programmes, eligibility and suitability for recognition of distinction, committee membership and external positions of influence.” (D5, 4.2)

#### Mentoring

Due to Athena SWAN, all departments have already established formal mentoring schemes for academic and research staff. Current actions aim to increase the uptake of mentoring, extend formal mentoring schemes to include all categories of staff and students, provide training in effective mentorship, and trial new approaches:

- “Keep encouraging postdocs and early career researchers to join the established [mentoring] scheme by publicising its benefits on our website and bulletin.” (D9, S16A)
- “Improve opportunities for mentoring, particularly for professional & support staff.” (D16, 3.6)
- “Provide training in effective mentorship to all managers.” (D12, 2.16)
- “Trial a scheme where junior clinical staff are assigned a senior sponsor who will be their advocate, including in the NHS clinical setting where the working environment can be challenging.” (D13, 1.15)

#### Support

Departments organise workshops, coffee mornings, peer support groups, and subject-specific events to help students, early career researches, as well as professional and support staff to make informed career choices. The main focus of support with career development of academic and research staff is on securing external research funding and establishing independence. Departments provide methodological training, administrative support, internal peer review, and help develop interview skills for fellowship and grant applications, commit internal funding and support of senior researchers to develop applications, and seek to increase teaching opportunities for junior researchers:

- “Provide a fellowship coordination process to ensure all applications receive the same support (e.g. internal review, mock interview).” (D13, 1.12)
- “Develop a mechanism for staff to ‘bid’ for funded protected time to work on fellowship and grant applications.” D8 5.2
- “Encourage senior staff to provide junior researchers with the opportunity to be a co-applicant on grant applications.” (D8, 5.3)
- “Generate greater teaching opportunities for junior researchers, and monitor gender balance of uptake.” (D15, 16)
- “Use Autumn School [for clinical medical students and foundation doctors] as a vehicle to inspire potential female academic psychiatrists.” (D14, 3.7)

### Flexible working and managing career breaks

#### Flexible and part-time working

Athena SWAN helps to improve arrangements for flexible and part-time working for all genders and groups of staff. Departmental action plans include interventions to promote the value of flexible and part-time working, raise awareness about the existing arrangements, formalise them through policies and guidelines, as well as extend to graduate research students:

- “Continue promote and de-stigmatise the value of flexible working and clarify the process for requesting this. Encourage culture of monitoring output rather than ‘presenteeism’.” (D10, 1.7)
- “Raise awareness of the flexible working policy: 26% of staff do not know about the flexible/part-time working policy.” (D2, 6.2)
- “Create guidelines for staff and their line managers explaining what part-time working entails and what to consider when deciding whether or not to become a part-time member of staff.” (D11, 8.6)
- “Explore opportunities for part-time work in [departmental] Clinical Research Facilities to facilitate career re-entry for clinicians.” (D13, 1.16)
- “Include part-time DPhil in graduate advertising, target clinical academic mentors to advertise the programme and explore sources of funding.” (D8, 2.3)

#### Managing career breaks

Athena SWAN has prompted departments to provide more support to women with managing maternity and other career breaks as well as introduce paternity and shared parental leave policies aimed at men. Departmental action plans contain further actions to improve the implementation of the existing policies and to commit resources to helping academic and research staff to return to research following a career break or a period of leave for caring responsibilities:

- “Plan how Shared Parental Leave will be managed in the department; also how to encourage women to consider sharing leave with their partner, and men to take leave.” (D5, 7.1)
- “Introduce ‘Buddy System’ for staff on maternity, paternity, caring or sick leave to help ensure that people are kept up-to-date with departmental decisions and policies.” (D2, 6.3)
- “Women returning to work after a period of maternity leave are to be given dispensation from teaching commitments.” (D9, S23a)
- “Continue to promote the Returning Carers’ Fund and encourage and support applications.” (D12, 5.1)

#### Childcare and family life

Departments work to enhance the provision of childcare in their specific locations and help staff to reconcile work and family life more broadly. Many departments improve information about available childcare services, invest into the provision of parking, breastfeeding facilities, and sponsored nursery places, as well as try to create a more family-friendly environment:

- “Provide pregnancy car parking space for expectant mothers who are finding their usual mode of transport to work challenging.” (D5, 7.6)
- “Our maternity/paternity focus group meeting raised the issue of a lack of breastfeeding support and facilities in the Department and the provision of a private room for breastfeeding is now underway.” (D4, S6.4)
- “Invest in sponsored nursery places.” (D2, 6.5)
- “Improve environment for women to discuss issues of home/ work-life balance with their line manager/supervisor” (D7, 11)
- “Support family friendly events in the divisions, to bring together staff, students and their families, and foster a sense of community in the department” (D13, 4.6)

### Organisation and culture

#### Embedding Athena SWAN into culture

Departments actively seek to embed the Athena SWAN principles into their culture by considering equality, diversity and inclusion as part of their values and identity, norms and procedures, social events, as well as working environments:

- “Design a set of core values to reflect the ethos of the Department; gain approval from [the Executive Committee]; publish on website.” (D12, 4.8)
- “Ensure diversity in imagery used on our website, in publicity materials and in social media.” (D16, 5.3)
- “Highlight Athena SWAN in all job advertisements.” (D10, 3.5)
- “Identify suitable speakers from outside the department to run a workshop for [group leaders] on how to create a positive and supportive culture in the lab.” (D12 2.14)
- “Make changes to working environments to improve the quality of working life.” (D3, 4.1)

#### Transparency and communications

Many actions to promote equality and inclusion focus on improving the transparency of departmental structures and decision-making, enabling equal access to key people and resources, and diversifying internal communication strategies:

- “Transparency, particularly regarding management structures and decision-making, will be further improved using a variety of communication strategies.” (D1, 17)
- “Set up a new sharepoint site for minutes of all meetings. Notify staff through the Weekly News that minutes have been published. Use multiple methods to give feedback on key issues including summarizing decisions in the department newsletter and at the termly department Open meeting.” (D11, 7.1)
- “Create admin postcards to distribute at admin surgeries to illustrate pipelines and contact persons for different processes (eg applying for a grant, recruiting a new staff member).” (D6, 5.2)
- “Ensure greater acknowledgement of success and achievement by adding success stories to the [Departmental] Digest and display screens in reception.” (D4, S5.10)

#### Zero tolerance of bullying and harassment

The Athena SWAN process provides departments with the opportunity to strengthen their core human resource policies to create a more positive culture for everyone. Most notably, all departments strive to eradicate bullying and harassment from the workplace by raising awareness of zero tolerance of bullying and harassment and providing resources to address it:

- “Reinforce messages about zero tolerance of bullying and harassment.” (D16, 5.2)
- “Initiate annual email to all postdocs reminding them of the bullying and harassment policy and the help available to raise awareness following the survey results.” (D5, 8.2)
- “We will include details of the Departmental bullying and harassment officers on the posters of key people and add this information to the intranet.” (D4, S5.7)
- “Appoint an external independent mediator/ listener to investigate the nature and extent of the problem (e.g. through targeted mini-survey)” (D13, 3.6)
- “Continue to provide a programme of in-house training on Bullying and Harassment.” (D8, 7.2)

#### Mandatory unconscious bias training

Another important human resources policy that many departments introduce as part of the Athena SWAN process is mandatory unconscious bias training. In addition to online training provided by the University, departments develop in-house online and in-person training and ensure completion by staff and students:

- “Introduce mandatory online equality & diversity and unconscious bias training for all staff and students.” (D13, 3.1)
- “Offer in-house unconscious bias training annually. Monitor compliance with compulsory training requirements. Include training records in staff database to check compulsory requirements.” (D6, 4.12)
- “We will chase up the 5% of group leaders who did not attend the unconscious bias training to complete an online version of the course to ensure 100% compliance.” (D5, 3.6)

#### Gender balance on committees

All departments work towards improving gender balance on committees, but with a varying degree of ambition. Whereas some department aim to improve gender balance relative to the proportion of group leaders, some other departments open up membership to all students and staff aiming to achieve absolute gender balance:

- “As number of female group leaders increases, increase their participation on committees. Aim to keep slightly ahead of simple proportion of female group leaders.” (D5, 6.2)
- “Review the membership of [Departmental] committees and identify more women as potential members (opening up membership of committees to students, [postdoctoral research assistants] and support staff where appropriate).” (D13, 3.4)
- “Ensure committees are gender-balanced. Monitor committee membership and attendance records. Rotate membership and chairs, with future vacancies appointed by advertisement and election. Monitor the reasons for requests to opt-out of committee membership.” (D15, 25).
- “Achieve gender balance on departmental committees.” (D14, 5.4)

#### Workload allocation model

The Athena SWAN process challenges departments to develop a fair and transparent workload allocation model, monitor it for gender bias, and use it for personal development review and promotion. Different departments are at different stages of implementing such a model, and in the majority of departments it remains limited to academic and research staff:

- “Set up a process, initially within PDR to look at workload, i.e., time spent on different core activities (research, internal administration and management committees, teaching and supervision, and outreach activities).” (D8, 4.3)
- “Work with [the Medical Sciences Division] to design a workload allocation model for clinical departments.” (D12, 4.9)
- “The workload model analysis will be continued on an annual basis to work towards parity between male and female group leaders. We will publicise our workload model within the University by arranging a workshop with other Departments in [Medical Sciences] and other Divisions to discuss best practice. This will also feed into refining our model.” (D1, 16)
- “Increase the proportion of staff who feel that their workload allocation is fair. Increase the transparency of workloads.” (D2, 5.4)

#### Timing of departmental meetings

As part of Athena SWAN bronze awards, the majority of departments have addressed the timing of departmental meetings and social events to make them possible to attend for staff with caring responsibilities and working part-time. Several departments, especially clinical ones, continue to reinforce the importance of inclusivity for all meetings and events:

- “Ensure that there is a high level of awareness of the concept of core hours by including it in staff induction material and in the monthly newsletter.” (D5, 7.8)
- “Promote inclusive meeting etiquette.” (D10, 5.2)
- “Schedule departmental meetings and seminars between 9.30am – 2.30pm wherever possible, and give considerable notice ahead of all day and evening events” (D13 4.3)
- “As far as practical, reduce scheduling of key meetings in school holidays.” (D5, 7.7)

#### Visibility of role models

All departments take action to promote visibility of role models and build gender equality and diversity into organisation of events and online materials:

- “Change the way the external seminar series is organised so that all potential seminar hosts have to nominate 2 speakers; 1 female and 1 male.” (D12, 4.3)
- “Increase the number of female research staff with a personal webpage.” (D4, S3.10)
- “Linking to the Staff Profiles area on the department website provide example career trajectories from [professional and support staff] at different points in their careers.” (D11, 5.1)
- “We are currently developing a website based on digital video interviews with university staff with a range of disabilities.” (D11, 10.2)

#### Outreach activities

The Athena SWAN process recognises the value of public engagement with science. This encourages departments to improve and reward outreach activities:

- “Improve profile of public engagement section of website and encourage people to submit examples of outreach to go on the News section and on display screens in reception.” (D4, S5.8)
- “Encourage more male researchers to take up Comms training and get involved in public engagement activities in the department – actively seek men to take part.” (D11, 8.3)
- “Run a series of public engagement/school outreach activities to attract female applicants from physical science disciplines, in which women are traditionally underrepresented.” (D12, 1.2)

### Complex contextually-embedded system of dynamic action planning

The five types of actions described above are organised into a complex contextually-embedded system of dynamic action planning and tailor-made to challenge gendered barriers to career progression at multiple, interacting levels (Figure 2). The systemic approach embraced herewith allows us to take a holistic view, considering that all parts of the system are interlinked. We thus consider the complex system in which the Athena SWAN scheme operates, acknowledging its non-linear character and the multitude of variables at play.

**Fig. 2.**
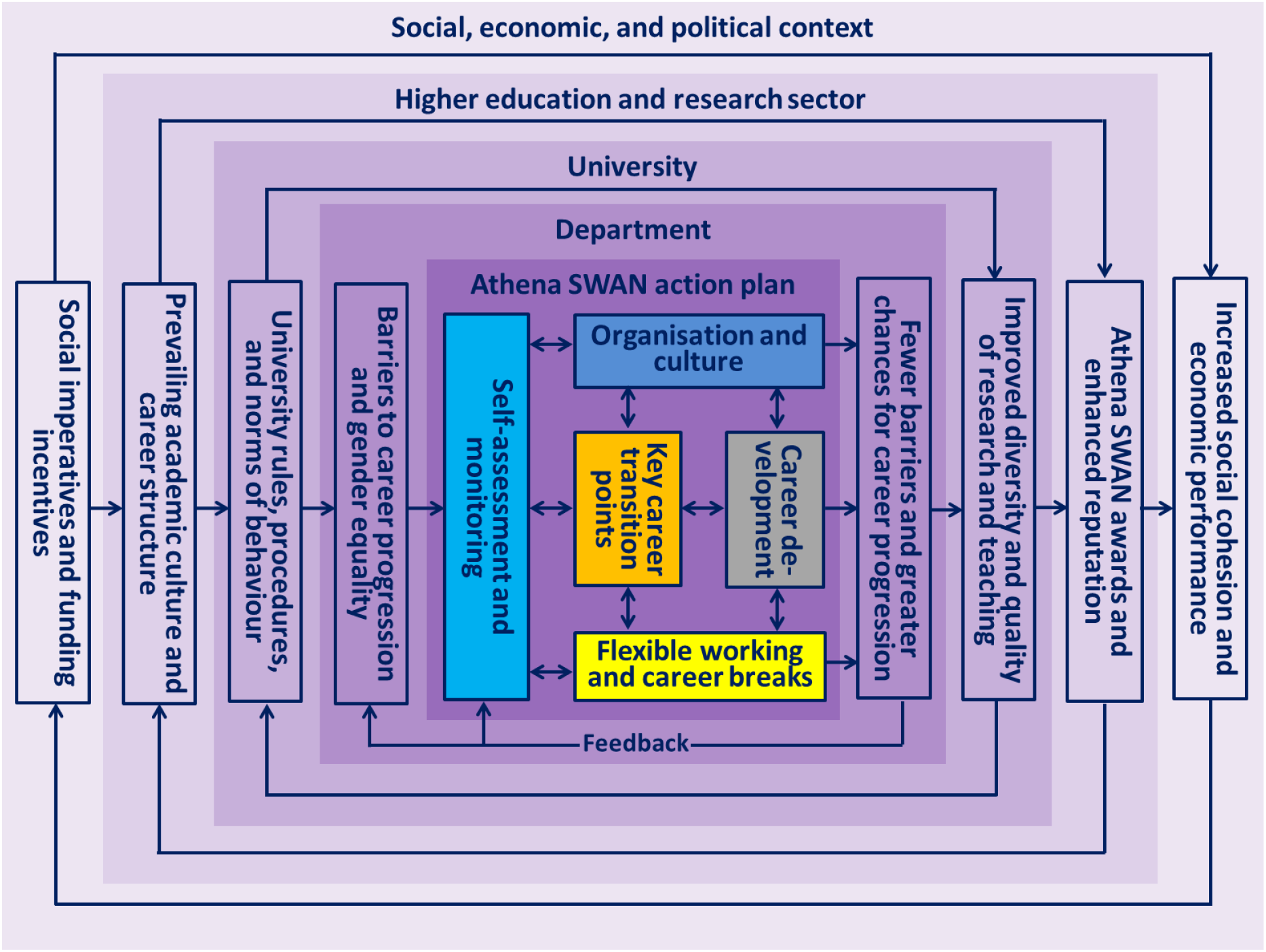
A complex contextually embedded system of Athena SWAN’s dynamic action planning.

Departmental action plans are dynamically linked to the wider social, economic, and political context, the higher education and research sector, and the university, which constitute a complex system. The widespread development and implementation of Athena SWAN Silver action plans resulted from the changes in the social, political, and economic context. Namely, the social progress imperative to improve gender equity in biomedical research prompted the government Department of Health to introduce NIHR funding incentives, in response to which the university and departments developed and implemented Athena SWAN Bronze action plans.

Following positive feedback in the form of an Athena SWAN Silver award, departments have developed and now implement Athena SWAN Silver action plans. They build on the outcomes of the previous Athena SWAN Bronze action plans, comments from the Athena SWAN peer review panel, and emerging best practice from the network of Athena SWAN Charter members and representing the wider higher education and research sector. The latter is particularly important because the prevailing academic culture and career structure influences the range of possibilities for change within individual universities. Hence, collective efforts of the entire higher education and research sector are often required to enable changes within individual universities. The context of individual universities is equally important because university rules, procedures, and both formal and informal norms of behaviour shape the range of interventions that departments can implement to remove barriers to career progression and gender equality within departments.

Interactions among the five types of actions in departmental action plans create a system-level organisation with positive feedback loops and new emergent properties. First, departmental self-assessment teams continuously assess data and evaluate the implementation of actions and their short- and medium-term impact. In response to the changing contextual factors and emerging evidence, they adapt on-going actions and develop new ones. In doing so, actions aimed at self-assessment and monitoring not only determine which actions are included in the action plan, but also regulate their implementation. In a complex system such as Athena SWAN, the agents of change are interconnected and affect each other. Thus, small alterations initiated by the self-assessment team can lead to large effects at a later point in time. This implies that increased probability of change is part of the expected impact of complex interventions.

Second, given that organisation and culture enable and constrain all interactions in the department, actions aimed at changing departmental organisation and culture also influence the other types of actions which adapt in response to the changing organisation and culture. There are multiple choices to make for staff and students that are subject to a range of contextual conditions, structural resistances and other constraints that impact career progression.

Third, actions aimed at flexible working and managing career breaks also dynamically interact with the other types of actions, in particular, with key career transition points and career development opportunities for staff and students taking advantages of flexible working arrangements and those taking career breaks. In complex systems, actions of different agents of change, such as individual choices of staff and students in terms of training activities, careers, courses etc. can lead to increased gender equality in the long run.

Finally, as more career development opportunities and better conditions for career progression through key career transition points emerge as a result of the implementation of action plans, key career transition points come faster and the chances of progressing to the next career stage in the new emergent conditions increase. Moreover, in contrast to complicated systems, complex systems are adaptive, which in this particular context means that they respond to the changes initiated through Athena SWAN. In this respect, action plans of Athena SWAN adapt to constantly moving targets and take into account new emergent conditions.

## Discussion

### Strengths and limitations in relation to previous research

Despite a growing body of literature on the design, implementation, and impact of gender equality interventions, there is a paucity of research on the complexity of gender equality interventions based on extensive empirical data [26]. To address this paucity of research, we embraced the complexity approach and analysed 16 departmental Athena SWAN Silver action plans in Medical Sciences at the University of Oxford, UK. To the best of our knowledge, it is the first study to show empirically that Athena SWAN is a complex social intervention and to discuss its implications for policy and practice. The study is based on the most extensive dataset of Athena SWAN interventions in Medical Sciences, but it is limited to a single site. Extending data collection and analysis to multiple sites is likely to capture a greater range of interventions and contextual factors. Another unique contribution of this study lays in comparing Athena SWAN Silver interventions with the EFFORTI typology of gender equality interventions in research and innovation in the wider European Research Area. Having a comparative European perspective can help generate insights on the strengths, limitations, and opportunities for further development of Athena SWAN. Given the current policy interest in introducing a gender equality award scheme similar to Athena SWAN in the wider European Research Area, our comparative analysis has a potential to inform policy and practice wide across Europe.

### Athena SWAN as a complex social intervention

The results presented above demonstrate that the Athena SWAN Silver action plans conform to some of the key considerations of complexity and thus can be usefully framed as complex social interventions embedded in a complex system.

First, addressing a specific area of gender inequality is not enough in complex systems when designing gender equality interventions, as a variety of interconnected factors are involved in the process [51]. Scholars [12, 52] show that the efficacy of gender equality interventions depends not only on the quality but also on the quantity of the measures implemented [53]. Complex systems are characterised by multiple actions and areas of intervention with a focus on the local dynamics [12]. Athena SWAN provides a dynamic system to address multiple cultural and structural aspects of gender inequality in accordance with the needs, baseline conditions, and emerging circumstances in participating departments. Athena SWAN’s focus on the local dynamics in departments is particularly important because departments rather than the central university make recruitment and promotion decisions, hold research funding, and provide working environments. Overall, 16 Medical Sciences departments implement 547 actions organised into five themes and 22 sub-themes. Actions are tailored to the specific departmental contexts and vary greatly in design, target populations, areas of intervention, and pace of implementation. Within departments, many actions are attuned to the context of different departmental divisions, institutes, centres, units, and research groups. Given that most of the departments are embedded in the context of hospitals and clinical facilities, actions vary between different medical specialties and basic science areas. Moreover, departments are often distributed across several campuses and physical locations, adding another layer of complexity to action planning.

Second, the complexity approach embraces the notion of the non-linearity of interventions and the constantly emerging conditions [12, 26, 54]. The non-linear relationship between inputs, outcomes and impact of gender equality action plans depend on the interaction of a variety of variables dynamically related to contextual factors. As Greenhalgh and Papoutsi [40] state, instead of a linear, cause-and-effect causality, the complexity paradigm is characterized by emergent causality, where manifold interacting features and “multiple uncertainties involved” produce effects that cannot be ascribed to one particular influence. Therefore, the design of complex gender equality interventions cannot afford to underestimate the inconsistency and unpredictability of the implementation of the actions [12]. Athena SWAN is not a stable arrangement, but a dynamic system that is in a continuous interaction with the environmental conditions, addressing new emerging conditions. Although initially Athena SWAN was set up to address barriers for women in academic and research roles, it has evolved to develop a broader focus on gender equality among all staff and students, taking into account considerations of intersectionality, such as sexuality, race, disability, age, and religion. Strikingly, the target population of the Athena SWAN Silver actions analysed by gender are predominantly all genders indiscriminately (87.9%) and nearly a half of the actions (48.4%) target all staff, students, or professional and support staff. It is likely that multiple uncertain components in the analysed action plans would create an additional layer of complexity during their implementation. Self-assessment teams continuously interact with the environmental conditions and address new emerging conditions. Namely, they monitor data and meet on a regular basis to map the outcomes of the action plans, evaluate feedback, and redesign actions to address new emerging conditions. Discussions and negotiations about different dimensions of change take place both internally within the departments and the university, as well as externally within the wider system constituted by the higher education and research sector, the social, economic, and political conditions, and the Athena SWAN community of practice.

Third, Rogers [37] highlights the challenges for impact assessment caused by the great number of variables involved in complex interventions, pointing out the persistent evolution of new variables in an adaptive system. In complex interventions, implementation, outcome, and impact become even less predictable, manageable and responsive to linear logic [55]. Therefore, the complexity approach focuses on the contribution – rather than attribution and causality – of interventions to the outcome and impact. Moreover, the complexity approach implies considering the increased probability of change as part of the desirable effect of complex interventions [12]. As a corollary, the expected impact of complex social interventions needs to be considered in terms of how they foster the conditions for change and increase the probability that change can occur in a particular complex setting [26, 56]. Athena SWAN action plans impact is hence expected in terms of contribution to change, improved conditions to foster change, and the increased probability that change can occur [26, 56].

The above discussed results provide examples of actions illustrating the wide range where impact can occur. The design, implementation, and impact of complex social interventions are best captured and assessed using a combination of quantitative and qualitative methods, case studies, and illustrative examples. Only a small proportion of actions, such as those regarding recruitment, promotion, and permanent contracts, aim to directly address the under-representation of women in certain positions. Therefore, the impact of only a small proportion of actions can be assessed using traditional quantitative indicators such as the number and proportion of women in certain positions. The majority of actions, especially those regarding organization and culture, career development, and flexible working and career breaks, aim to improve conditions to foster change and increase the probability that change can occur. Moreover, a large number of actions regarding self-assessment and monitoring may also create emergent effects, such as the Hawthorne effect whereby staff modify their behaviour in response to the awareness of being observed. Therefore, assessing the impact of the majority of actions would require a combination of quantitative and qualitative methods taking into account possible emergent effects.

### Practical implications for impact assessment

While the assessment of the impact of the analysed Athena SWAN Silver actions is beyond the scope of this paper, the preceding discussion on Athena SWAN as a complex social intervention presents us with three practical implications for impact assessment^4^.

First, widening the areas where impact can be recognized. This requires going beyond what traditionally is perceived as impact and often measured quantitatively to also include qualitative parameters. Likewise, this requires taking into consideration the *ability* of interventions to produce impact within the targeted areas. Thus, it is important to anticipate the ability of the design and implementation of certain interventions to foster the expected “conditions for change” linking design, implementation and effect of interventions to adequate conditions to produce impact [56].

Second, identifying potential impacts on the creation of the right conditions for change in a medium- and long-term perspective. This probabilistic stance makes impact assessment of complex interventions “less deterministic and more substantive” [12]. Probability can then be assessed “through a set of indicators pointing to the activation of internal change processes (e.g. the successful involvement of internal and external actors, the modification of relevant rules, and the creation of internal groups of actors aimed at pursuing change), as internal processes are likely to produce additional impacts with time” [26].

Third, accounting for the influence of the context, the local dynamics, and the emerging conditions [57, 58]. Complex systems constantly adapt, not least, in response to interventions [36]. What seems to be a dominant cause of inequality at one point in time might shift later due to the constant interplay of a multitude of contextual factors [26]. As Greenhalgh and Papoutsi [40] state “we need to develop capability and capacity to handle the unknown, the uncertain, the unpredictable and the emergent” [59].

### Strategic opportunities to strengthen gender equality policies in the European Research Area

A comparison of the SWAN Silver gender equality action plans with the EFFORTI typology of gender equality interventions in research and innovation shows that Athena SWAN is the single most comprehensive and inclusive gender equality scheme in Europe. Athena SWAN covers approximately three quarters of gender equality intervention types used in the wider European Research Area. While gender equality interventions in the European Research Area tend to focus primarily on women in academic and research roles, Athena SWAN has a broader focus on all categories of staff and students, predominantly, regardless of their gender, taking into account considerations of intersectionality, such as sexuality, race, disability, age, and religion.

Athena SWAN also has a more contextually-embedded, country-wide systemic approach to action planning than any other single gender equality scheme in Europe, especially, with regard to system-level interventions related to institutionalised self-assessment teams, considerations of intersectionality, key career transition points, career development, mandatory training on unconscious bias and bullying and harassment, timing of meeting and events, and a workload allocation model. Gender equality policies in the wider European Research Area can benefit from exploring Athena SWAN’s contextually-embedded systemic approach to dynamic action planning and inclusive focus on all genders and categories of staff and students. Yet, Athena SWAN has two limitations with regard to intervention types. Whereas some European countries intervene to introduce quotas, positions, funding, and single-sex degree and specialisation courses, Athena SWAN does not promote such interventions because, under the UK Equality Act 2010, they may be interpreted as positive discrimination and therefore deemed unlawful. Moreover, Athena SWAN misses the opportunity to promote the integration of sex and gender dimension in research and education, which is particularly important both in the wider European Research Area and globally.

Together with fostering gender balance in research teams and in decision-making, integrating the gender dimension in research and education is one of three key objectives for promoting gender equality in research and innovation in Europe [23]. Research shows that increasing the participation of women in research and innovation “will not be successful without restructuring institutions and incorporating gender analysis into research” [60, 61]. Grounded on the Stanford University project “Gendered Innovations”^5^, the EC report “*Gendered Innovations: How Gender Analysis Contributes to Research”* [23] demonstrated how sex and gender analysis enhances the scientific quality, societal relevance, and business value of research and provided tools and guidance to do so^6^.

Promoting the integration of sex and gender analysis in research and education represents a strategic opportunity to strengthen Athena SWAN in the given research-intensive study setting. There is a growing body of evidence on how the incorporation of sex and gender in research leads to a better health care [62] or how the disregard of gender aspects [63–65] leads to sub-optimal, sometimes harmful health care [32, 60, 66]. The world’s leading health research funder, the US National Institutes of Health Research (NIH), has made it mandatory in 2016 that all researchers account for sex as a biological variable [67]. Many other health research funders world-wide have also introduced policies that require all grant applicants consider sex and gender variables in research design [68]. Likewise, The European Association of Science Editors has introduced the Sex and Gender Equity in Research (SAGER) guidelines to maximise the generalisability and applicability of research findings to clinical practice [69]. The SAGER guidelines help editors and researchers to ensure the adequate reporting of sex and gender information in study design, data analysis, results, and interpretations of findings [70, 71]. Moreover, including sex and gender analysis in the curricula of Medical Sciences courses helps students improve their study design, analysis, and reporting skills [60], and gender-sensitive curricula, portraying gender in a non-stereotypical way, may make academic and research careers in Medical Sciences more attractive to all irrespective of gender [72].

## Conclusions

To activate effective gender equality structural and cultural change, it is necessary to acknowledge and operationalize the notion of complexity as a frame of reference. Athena SWAN is the single most comprehensive and inclusive gender equality scheme in Europe and can be strengthened further by promoting the integration of sex and gender analysis in research and education. Gender equality policies in the wider European Research Area can benefit from exploring Athena SWAN’s contextually-embedded systemic approach to action planning and inclusive focus on all genders and categories of staff and students.

## Abbreviations

EC: European Commission
EFFORTI: Evaluation Framework for Promoting Gender Equality in Research and Innovation
ERA: European Research Area
GEAR: Gender Equality in Academia and Research
GEDII: Gender Diversity Impact – Improving research and innovation through gender diversity
Gender-NET: Promoting Gender Equality in Research Institutions and Integration of the Gender Dimension in Research Content
GENERA: Gender Equality Network in the European Research Area
NIHR: National Institute for Health Research
NHS: National Health Service
PRAGES: Practicing Gender Equality in Science
R&I: Research and Innovation
RRI: Responsible Research and Innovation
RTD: Research, Technology and Development
SAT: Self-assessment team
STAGES: Structural Transformation to Achieve Gender Equality in Science
STEM: Science, Technology, Engineering and Mathematics
SWAFS: Science With and For Society
UK: United Kingdom
UN: United Nations

## Declarations

### Acknowledgements

We thank Professor Trish Greenhalgh, Department of Primary Health Care Sciences, University of Oxford for comments and suggestions on an earlier version of the manuscript.

### Availability of data and materials

The datasets analysed during the current study are publicly available online at: https://www.admin.ox.ac.uk/eop/gender/athenaswan/applications/

### Authors’ contributions

EKS structured and led the drafting of the manuscript, including the introduction, methods, results, discussion, and conclusions. EKS and PVO conceived and designed the study, and analysed the data. PVO led data collection and analysis and co-drafted parts of the introduction, methods, results, discussion, and conclusions. VK and LRH analysed actions against the relevant sections and sub-sections of the 2015 Athena SWAN Charter Awards Handbook and drafted parts of the results sections. All authors have read and approved the final version.

### Ethics approval and consent to participate

Not applicable. A wider programme of research on the activities of the NIHR Oxford Biomedical Research Centre from 2017 to 2022 received ethics clearance through the University of Oxford Central University Research Ethics Committee (R51801/ RE001).

### Consent for publication

Not applicable.

### Authors’ information

**EKS** is Associate Professor and Research Director at the Department of Political Science, Aarhus University. She specializes in science and innovation policy; Responsible Research and Innovation; gender in knowledge production and research organizations; European gender policies and strategies; impact of knowledge governance and policy interventions; research evaluation; European research policy and governance. She has been involved in a number of projects funded by the EU and has been frequently engaged by the European Commission as expert in the evaluations of FP6, FP7 and Horizon 2020 project proposals. She is the European Commission appointed Danish expert member of the European RTD Evaluation Network and expert member of the Horizon 2020 Advisory Group for Gender.

**PVO** is Senior Research Fellow in Health Policy and Management in the Radcliffe Department of Medicine, University of Oxford. He is leading a multi-disciplinary programme of research and policy advocacy on gender equity across medical and social sciences. His current research includes building an evidence base to accelerate women’s advancement and leadership through systematic reviews; developing markers of achievement, metrics, and indicators for assessing and monitoring gender equity; investigating the gender dimension of responsible research and innovation to maximise scientific, societal and economic returns on investment in research; and examining ways of creating an inclusive university culture.

**LRH** is a visiting academic at the Radcliffe Department of Medicine. Her research has built an evidence base in NHS patient experiences research and influenced governmental healthcare policy. Her current research involves a Horizon 2020 European research project on developing markers for achievement concerning Gender in Biomedical research centres and gender equity and responsible research and innovation.

**VK** is the Chief Operating Officer at the NIHR Oxford Biomedical Research Centre (BRC). VK is a PhD holder in Cardiovascular Physiology and Genetics and holds an Executive MBA from the Said Business School, Oxford University. VK is also a Senior Research Fellow at the Department of Primary Health Care Science, University of Oxford. Her research interests focus on public health issues such as smoking, drug addiction and loneliness.

## Footnotes

1 RRI is promoted through actions on thematic elements (public engagement, gender, open access, ethics, science education), as well as integrated actions that address institutional change to foster the uptake of the RRI approach by stakeholders and European organisations. For a more detailed description of the RRI concept see European Commission. Responsible Research and Innovation. Europe’s ability to respond to societal challenges. European Union. 2012, Brussels.

2 See the GEAR tool https://eige.europa.eu/gender-mainstreaming/toolkits/gear

3 An interesting UK study containing 50 potential interventions representing good practice or positive action, and addressing cultural, organisational and individual barriers to gender equality, ranked by participants according to their perception of priority, is presented in a typology developed by Bryant et al. 49. Bryant LD, Burkinshaw P, House AO, West RM, Ward V. Good practice or positive action? Using Q methodology to identify competing views on improving gender equality in academic medicine. BMJ Open. 2017;7(8).

4 The strategies are developed from Kalpazidou Schmidt and Cacace 2017.

5 See http://genderedinnovations.stanford.edu/

6 Practical methods of sex and gender analysis for researchers have been developed and checklists have been provided by experts. The Gendered Innovations project has for instance developed practical examples of how sex and gender analysis leads to gendered innovations in Science, Health & Medicine, Engineering and the Environment, see http://genderedinnovations.stanford.edu/

