## Additional file 1 for "Understanding the Athena SWAN award scheme for gender equality as a complex social intervention in a complex system: analysis of Silver award action plans in a comparative European perspective"

**Additional file 1: Overview of the intervention typology developed in the EFFORTI project**

| **Type of intervention** | **Intervention format** |
| --- | --- |
| 1. **Policies** | 1. Mainstreaming actions 2. Gender equality/action plan 3. Gender budgeting |
| 2. **Non-discrimination** | 4. Gender-sensitive practices for the attribution of tasks 5. Gender-sensitive study and working conditions (e.g. alternative study plans for pregnancy during laboratory work period) 6. Gender-sensitive Human Resources management 7. Guidelines regarding gender specifics |
| 3. **Composition & Integration** | 8. Definition of targets regarding gender balance in decision-making positions 9. Definition of targets regarding gender balance in research groups 10. Institution of quotas |
| 4. **Advancement** | 11. Mentoring programmes 12. Gender-sensitive practices for assessment 13. Introduction of chairs and positions reserved to women 14. Support to career development (counselling) 15. Empowerment schemes |
| 5. **Recruitment** | 16. 16. Campaigns for inspiring women for Mathematics, information technology, natural sciences and technology (MINT) subjects |
| 6. **Monitoring** | 17. Monitoring appointments, promotions, or attributions of tasks |
| 7. **Deconstructing Excellence** | 18. Revision of internal policies regarding promotions 19. Revision of internal policies regarding staff appointments |
| 8. **Gender Awareness & Bias** | 20. Training courses (different targets) |
| 9. **Leadership Accountability** | 21. Implementation of gender-sensitive leadership and personnel development |
| 10. **Funding** | 22. Targeting funding practices to improve women’s access to research funding 23. (Targeted) funding to improve the integration of gender dimension in research 24. Targeted funding practices to encourage research organisations to promote gender equality measures 25. Special funding for women researchers |
| 11. **Research** | 26. Gendered user involvement 27. Inclusion and monitoring the integration of the gender dimension and impact |
| 12. **Knowledge** | 28. Dissemination of information material 29. Revision of teaching curricula and texts 30. Introduction of single-sex degree and specialisation courses 31. Provision of gender and women studies or modules 32. Integrating the gender dimension in tertiary education |
| 13. **Visibility** | 33. Networking 34. Activities to make women (and their research) visible (e.g. introduction of awards reserved for women) 35. Role models |
| 14. **Care & Family Life** | 36. Support in period of absence for family needs 37. Schemes for women returners 38. Care services and facilities (for children, the elderly, and others) 39. Support to mobility, including spouse relocation schemes |
| 15. **Work-Life Balance** | 40. Introduction of flexible working hours |
