## Additional file 2 for "Understanding the Athena SWAN award scheme for gender equality as a complex social intervention in a complex system: analysis of Silver award action plans in a comparative European perspective"

**Additional file 2: Departmental codes**

| **Code** | **Name of department** | **Date of Athena SWAN award application** |
| --- | --- | --- |
| **D1** | Department of Biochemistry | April, 2015 |
| **D2** | Department of Experimental Psychology | April, 2015 |
| **D3** | Department of Paediatrics | April, 2015 |
| **D4** | Department of Physiology, Anatomy and Genetics | April, 2015 |
| **D5** | Sir William Dunn School of Pathology | April, 2015 |
| **D6** | Nuffield Department of Clinical Neurosciences | April, 2015 |
| **D7** | Nuffield Department of Obstetrics & Gynaecology (as of 2017, Nuffield Department of Women's & Reproductive Health) | April, 2015 |
| **D8** | Nuffield Department of Population Health | April, 2015 |
| **D9** | Nuffield Department of Orthopaedics, Rheumatology and Musculoskeletal Sciences | April, 2015 |
| **D10** | Nuffield Department of Clinical Medicine | November, 2015 |
| **D11** | Nuffield Department of Primary Care Health Sciences | April, 2017 |
| **D12** | Department of Oncology | April, 2016 |
| **D13** | Radcliffe Department of Medicine | November, 2015 |
| **D14** | Department of Psychiatry | November, 2014 |
| **D15** | Nuffield Department of Surgical Sciences | April, 2016 |
| **D16** | National Perinatal Epidemiology Unit | November, 2016 |
